# The effect of domain growth on spatial correlations

**DOI:** 10.1101/041491

**Authors:** Robert J. H. Ross, C. A. Yates, R. E. Baker

## Abstract

Mathematical models describing cell movement and proliferation are important research tools for the understanding of many biological processes. In this work we present methods to include the effects of domain growth on the evolution of spatial correlations between agent locations in a continuum approximation of a one-dimensional lattice-based model of cell motility and proliferation. This is important as the inclusion of spatial correlations in continuum models of cell motility and proliferation without domain growth has previously been shown to be essential for their accuracy in certain scenarios. We include the effect of spatial correlations by deriving a system of ordinary differential equations that describe the expected evolution of individual and pair density functions for agents on a growing domain. We then demonstrate how to simplify this system of ordinary differential equations by using an appropriate approximation. This simplification allows domain growth to be included in models describing the evolution of spatial correlations between agents in a tractable manner.

## 1 Introduction

Many important biological processes during development involve the movement and proliferation of cell populations on growing domains [1]. For example, cranial neural crest stem cells, a subset of a migratory cell population that give rise to a diverse lineage, have been shown to migrate along the developing cranofacial region in embryonic chickens [2-4]. Similarly melanoblasts, neural crest precursors to melanocytes, have been shown to migrate through the developing dorsal lateral epithelium in the embryonic mouse [5-7].

In both of the aforementioned examples, individual-based models (IBMs) have played an important role in research into these cell migratory processes [3, 4]. Studies involving IBMs have shown, in the case of melanoblasts, that the distribution of the migrating cells is thought to determine fur patterning and pigmentation defects such as piebaldism [8]. In the case of cranial neural crest stem cells, IBMs have helped to elucidate the mechanisms by which a cell becomes a ‘leader’ or a ‘follower’ in the collective cell migration process [2-4]. IBMs allow an intuitive representation of cells (referred to as ‘agents’ in the IBM), and allow for complex behaviours, such as cell-cell interactions and volume exclusion, to be easily assigned to agents in the model [9-12]. Importantly, IBMs can capture the effects of spatial correlations and heterogeneity in agent populations, and the ramifications spatial correlations can have on density-dependent processes such as cell migration and proliferation [13-18].

IBMs are also often amenable to approximation by population-level continuum models. Accurate continuum approximations of IBMs are important tools for understanding biological systems as, in contrast to IBMs, they generally allow for more mathematical analysis. This analysis can be crucial to form a mechanistic understanding of biological systems, which is not always apparent (or feasible) from simply studying the averaged results of a large number of repeats of an IBM. For instance, the exploration of a large parameter space. However, in certain scenarios standard mean-field partial differential equation (PDE) descriptions of IBMs, such as those describing the expected evolution of the population density, suffer from the limitation that they neglect to incorporate the impact of spatial correlations and clustering. Therefore, in order to derive accurate continuum approximations of IBMs it is often necessary to include the effects of spatial correlations in continuum models [13-23]. Furthermore, having the mathematical tools to directly compute spatial correlations allows them to be analysed, which can give important insights into the biological process being studied. For instance, spatial correlations indicative of different types of cell-cell interactions can be observed in cell populations [24-26], and spatial correlations between cells are thought to play an important role in tumour growth [27].

In this work we examine how domain growth affects the evolution of individual and pair density functions for agents in an IBM. A large body of literature already exists concerning the evolution of individual and pair density functions on static domains [13-18], the most striking examples of which show that standard mean-field PDE descriptions can be wholly insufficient approximations of the evolution of the agent density in IBMs in certain scenarios [13, 16]. We therefore also display how to integrate the results presented here into pre-existing models. In doing so we simplify the implementation of the methods we present so that they can be more easily applied to the study of complex systems.

The outline of this work is as follows: to begin we introduce our one-dimensional IBM and domain growth mechanism in Section 2.1. We then define the individual and pair density functions, and derive a system of ordinary differential equations (ODEs) describing the evolution of the individual and pair density functions with respect to time on a growing domain in Section 2.2. To test the accuracy of this system of ODEs we compare its numerical solution with ensemble averages of the individual and pair agent densities from the IBM for a range of initial conditions and parameter values in Section 3. In Section 4 we integrate domain growth into existing models for calculating the evolution of pairwise spatial correlations. These models are typically used to correct mean-field approximations for the evolution of the agent density in an IBM by taking spatial correlations into account. In Section 5 we conclude with a discussion of the results presented.

## 2 Model

In this section we first introduce the IBM and the domain growth mechanism we employ throughout this work. We then introduce the individual and pair density functions and derive a system of ODEs describing the evolution of these functions in the IBM.

### 2.1 One-dimensional IBM and the domain growth mechanism

We use an agent-based, discrete random-walk model on a one-dimensional regular lattice with lattice spacing Δ [28] and length *L(t)*, where *L(t)* is an integer describing the number of lattice sites. Throughout this work the lattice site spacing, Δ, is always equal to one. All simulations are performed with periodic boundary conditions. Each agent is assigned to a lattice site, from which it can move or proliferate into an adjacent site. If an agent attempts to move into a site that is already occupied, the movement event is aborted. Similarly, if an agent attempts to proliferate into a site that is already occupied, the proliferation event is aborted. This process, whereby only one agent is allowed per site, is referred to as an exclusion process [28]. Time is evolved continuously, in accordance with the Gillespie algorithm [29], such that movement, proliferation and growth events are modelled as exponentially distributed ‘reaction events’ in a Markov chain. Attempted agent movement or proliferation events occur with rates *P*_*m*_ or *P*_*p*_ per unit time, respectively. That is, *P*_*m*_*δt* is the probability of an agent attempting to move in the next infinitesimally small time interval *δt*. Throughout this work the initial agent distribution for all simulations is achieved by populating lattice sites uniformly at random until the required density is achieved.

We employ a stochastic growth mechanism in one dimension in which the insertion of new lattice sites into the domain occurs with rate *P*_*g*_*L(t)* per unit time. That is, each individual lattice site undergoes a growth event with rate *P*_*g*_ > 0. When a growth event occurs a site on the domain is selected uniformly at random and a new (empty) lattice site is inserted at the selected site, as can be seen in Fig. 1. For notational convenience from this point on we write *L(t)*as *L*.

Figure 1 also displays how the distance ‘r’ between two lattice sites on the domain is defined. This distance *r* is measured between the centres of the lattice sites and in the clockwise direction. This means that on a domain of length *L* there are *L* counts of each distance *r* ∈ [1,…, *L* − 1] between lattice sites.

**Figure 1:**
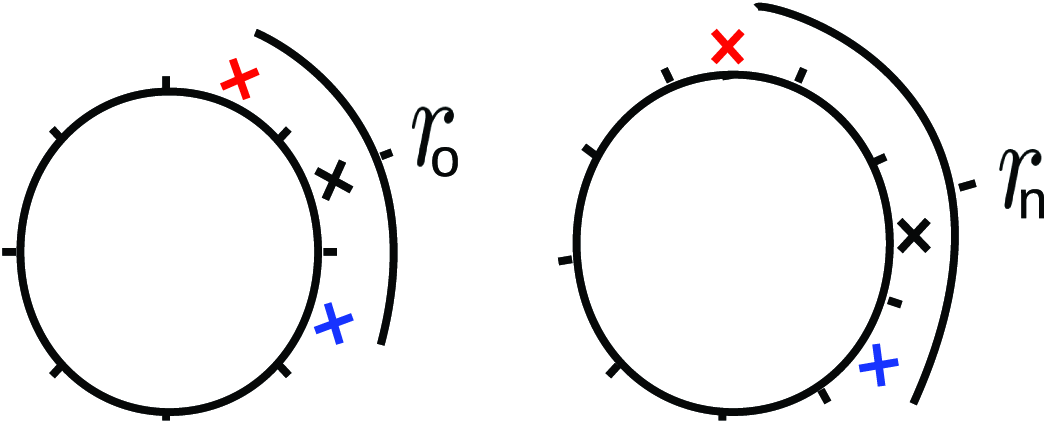
(Colour online). A one-dimensional domain with periodic boundary conditions can be represented as a circle. A site marked by a black cross has been chosen to undergo a growth event. Following this event the site marked by the black cross and its contents are moved one lattice spacing in the clockwise direction on the circle and a new lattice site is inserted. The new lattice site is empty. The distance between the lattice site marked by the red cross and that marked by the blue cross is measured in the clockwise direction. In this example the distance between the two sites initially, *r*_0_, is 2. Following the growth event the distance between these lattice sites, *r*_n_, is 3.

### 2.2 System of ODEs

We begin by introducing the density functions. A density function, 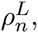, is defined as the probability that *n* sites have given occupancies when there are *L* sites in the domain [13-18]. For instance, a pairwise density function, 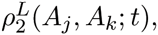 is the probability that sites *j* and *k* are both occupied by an agent ‘A’ at time *t* on a domain of length *L* (i.e. contains *L* sites). An individual density function, 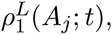 is the probability that site *j* is occupied by an agent *‘A’* at time *t* on a domain of length *L*.

In the course of this derivation the pairwise density functions will be rewritten in terms of the distance between two sites, *r*. We can do this as we assume translational invariance of the density functions throughout this work. We are able to assume translational invariance because the initial condition used in our IBM simulations is achieved by populating sites uniformly at random until we have achieved a required initial density. Two consequences of assuming translational invariance are

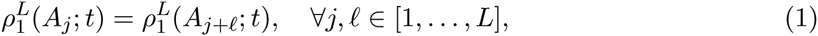

and

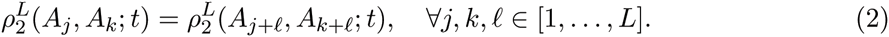

Equation (1) simply means that the probability of a site being occupied is independent of its location on the domain. Similarly, Eq. (2) means that the probability of two sites a distance *r* apart being occupied is independent of the location of the two sites on the domain (i.e *j* + *r* = *k*). Importantly, Eqs. (1) and (2) allow us to simplify the following derivation greatly as 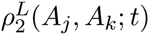 can be written as a function of the distance between two sites, so that

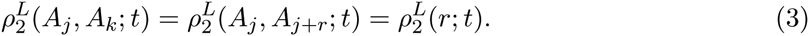

This ‘abuse’ of notation, whereby 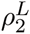 is rewritten as a function of the distance between two lattice sites as opposed to the lattice sites themselves, will prove useful in the following derivation.

To begin with, we only model the effects of domain growth on the density functions (the effects of agent motility and proliferation on the density functions are included later). On a growing domain the individual probability density functions evolve according to

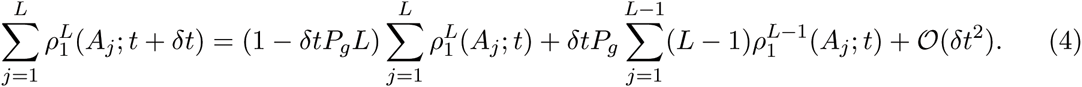

That is, the sum of individual probability density functions on a domain of length *L* at [*t* + *δt*) corresponds to the following terms on the right-hand-side (RHS) of Eq. (4): i) the sum of individual probability density functions at time *t* multiplied by the probability that no growth event occurs in [*t, t* + *δt*); and ii) the sum of individual probability density functions at time *t* on a domain of length *L −* 1 multiplied by the probability that a growth event occurs. To simplify Eq. (4) we rewrite 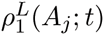 as *c*^*L*^*(t)* to obtain

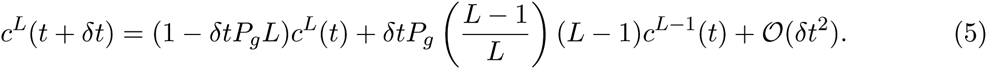

Finally, if we rearrange Eq. (5) and take the limit as *δt* → 0 we arrive at

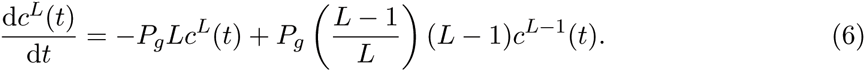

Equation (6) is a system of coupled ODEs for the evolution of the individual agent density for each domain length *L*. That is, the probability of a site being occupied on a domain of length *L* at time *t*.

We now consider the evolution of the pairwise density functions. As previously discussed we can rewrite the pairwise density functions 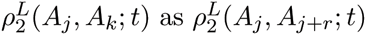for *r* = 0,…, *L* − 1 (if *j + r* > *L* the pairwise density function is instead 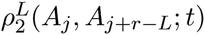 as the domain boundary is periodic). The evolution of the pairwise density functions on a growing domain can be written
as

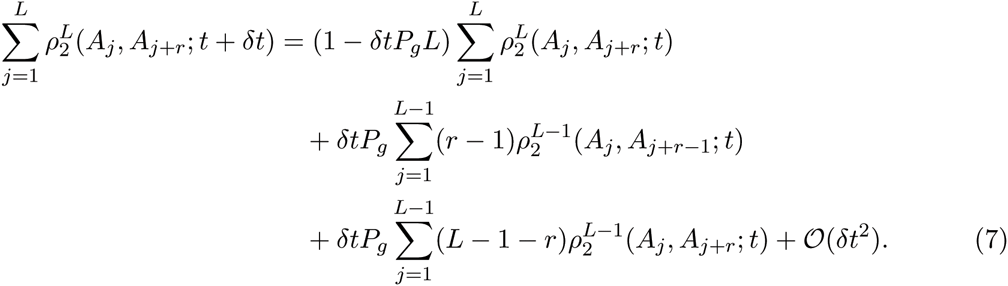

The left-hand-side (LHS) of Eq. (7) is the sum of pairwise density functions for lattice sites a distance *r* apart on a domain of length *L* at [*t* + *δt*). The RHS of Eq. (7) corresponds to: (i) the sum of pairwise density functions for lattice sites a distance *r* apart on a domain of length *L* at time *t* multiplied by the probability that no growth event occurs in [*t, t* + *δt*); (ii) the sum of pairwise density functions for lattice sites a distance *r −* 1 apart on a domain of length *L −* 1 multiplied by the probability that a domain growth event occurs in the interval between these sites (i.e. in one of *r* − 1 sites), thereby moving two lattice sites to a distance *r* apart on a domain of length *L* at time [*t* + *δt*); (iii) the sum of pairwise density functions for lattice sites a distance *r* apart on a domain of length *L −* 1 multiplied by the probability that a growth event does not occur in the interval of length *r* between the sites (i.e. it occurs in one of *L −* 1 − *r* sites instead), therefore moving two lattice sites a distance *r* apart on a domain of length *L −* 1 into a domain of length *L* at [*t* + *δt*).

To simplify Eq. (7) we can rewrite 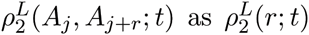 as in Eq. (3). If we substitute 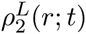 into Eq. (7) we obtain

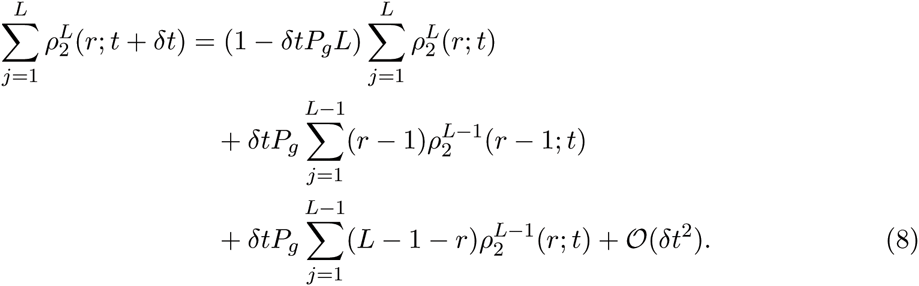

Equation (8) can be simplified and rewritten as

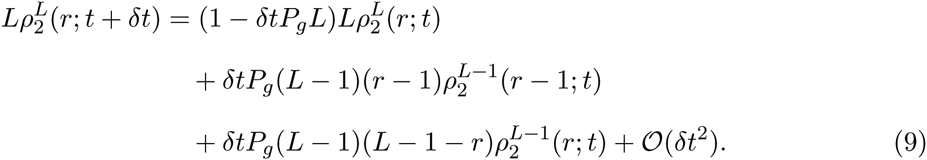

Finally, if we rearrange Eq. (9) and take the limit as *δt* → 0 we arrive at

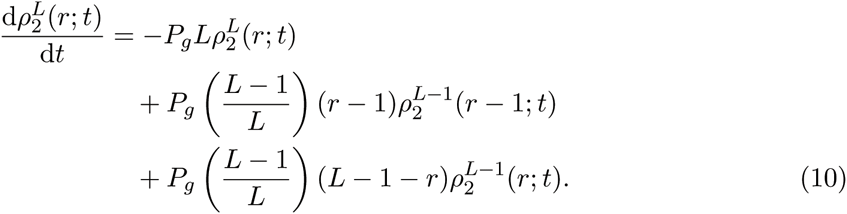

Equation (10) describes the evolution of all pairwise density functions for distances *r* = 1,…, *L*− 1 on a growing domain. Similar to Eq. (6), Eq. (10) constitutes a system of coupled ODEs for the evolution of the pairwise density functions at a distance *r* for each domain length *L*.

### 2.3 Motility

It has previously been shown how to include the effect of agent motility in Eq. (10), albeit for the non-growing case. However, as the derivation is similar we refer the reader to Appendix A for the derivation and simply state the result here [13]. In the case that *r* = 1 the evolution of pairwise density functions for motile agents on a growing domain is given by

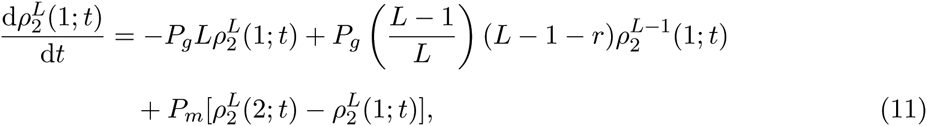

and for *r* > 1

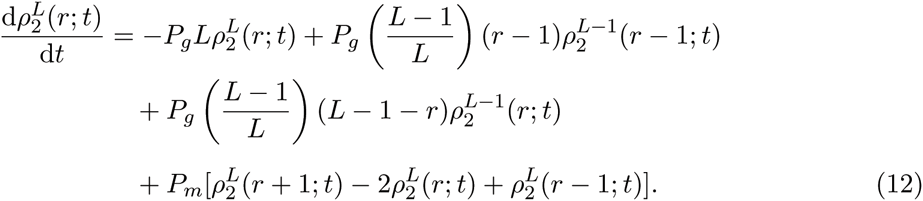

The addition of agent motility does not affect the evolution of individual density functions given by Eq. (6) [13].

## 3 Results

We now present results for motile agents on a growing domain, in which domain growth is exponential. The lattice state of site *j* in the *i*th realisation of the one-dimensional IBM is described by variable *σ*_*ji*_ (i.e. occupied by an agent or unoccupied). This means the normalised average agent density for the ith realisation in the one-dimensional IBM is

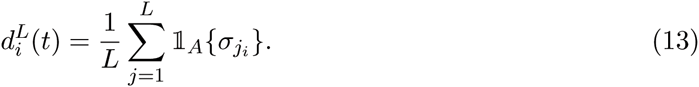

Here 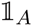 is the indicator function for the occupancy of a lattice site (i.e. 1 if an agent occupies lattice site *j*, and 0 if it does not). If we take the ensemble average of Eq. (13) over many repeats we obtain the average density. Given our initial conditions, the expected (average) probability of two sites a distance *r* apart being occupied can be calculated exactly as:

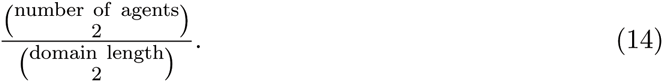

That is, the number of agents on the domain choose two, divided by the number of lattice sites choose two. For an initial condition in which all lattice sites are occupied this is simply unity, and for an initial condition containing only one agent it is undefined (because at least two agents are required for a pair density). Therefore, Eq. (14) provides the means of calculating the initial values for Eqs. (11) and (12). The initial condition for the individual density functions, Eq. (6), is simply the initial density at the initial domain length *L*_0_, and zero for all other lengths (i.e. for *L* > *L*_0_). That is

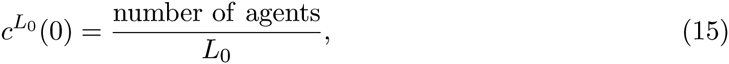

and

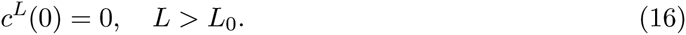

We calculate the pairwise counts between agents from IBM simulations in the following way: at each time-point every distance between an agent and all other agents is recorded. For a one-dimensional domain with periodic boundaries there are two distances between a pair of agents, of which we record the shorter distance. This count, defined as *s*(*r, L, t*), is then normalised so that

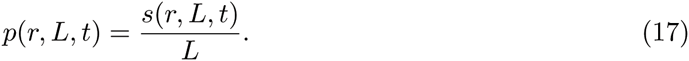

To make comparisons between ensemble averages from our discrete model and numerical solutions of the system of ODEs straightforward we only solve the system of ODEs given by Eqs. (11) and (12) for distance

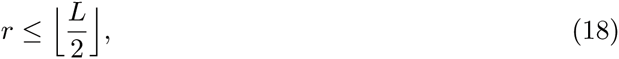

where ⌊·⌋ represents the floor function. Implementing Eq. (18) makes the results directly comparable to much of the previously published work on density functions, whereby the shortest distance between two sites is taken as the distance between these two sites [24-26]. In cases where 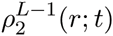 does not exist due to the condition imposed by Eq. (18), for instance when the domain length is even, we set 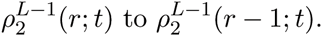 This is simply the anti-clockwise distance between the same two sites. These two sites must have the same joint probability of occupancy independent of the direction taken to measure the distance between them. In cases where 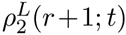 does not exist due to the condition imposed by Eq. (18), it is set to 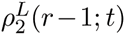 if the domain length *L* is even (again, by considering the symmetry of the domain), and 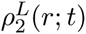 if the domain length *L* is odd.

Finally, given we are modelling domain growth it is necessary to truncate our state space, and so we truncate the derived ODE model at approximately three times the expected final domain length. As the domain grows exponentially the expected final domain length is simply

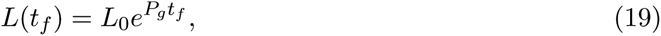

where *t*_*f*_ is the end time-point of the simulation. The truncation length is represented by *L*_*trunc*_ in the figures. Small alterations to *L*_*trunc*_ had no noticeable effect on the results presented here. We solve Eqs. (6), (11) and (12) using MATLAB’s `ode15s`.

### 3.1 Motile agents on a growing domain

In Fig. 2 the evolution of the individual density functions can be seen for two different simulations with different parameters. In Fig. 2 (a) results are shown for a domain with an initial length of ten lattice sites and ten agents. In Fig. 2 (b) results are shown for a domain with an initial length of thirty lattice sites and fifteen agents. We see an excellent agreement between the numerical solution of Eq. (6) and the ensemble averages from the IBM, Eq. (13), for all domain lengths for both simulations. This is to be expected as Eqs. (6) and (11), (12) are all exact. The probability of a site being occupied with the domain at a given length increases and then decreases for all domain lengths (apart from the initial domain length *L*_0_). This is because initially all domains are of length *L*_0_ (see Eq. (16)).

**Figure 2:**
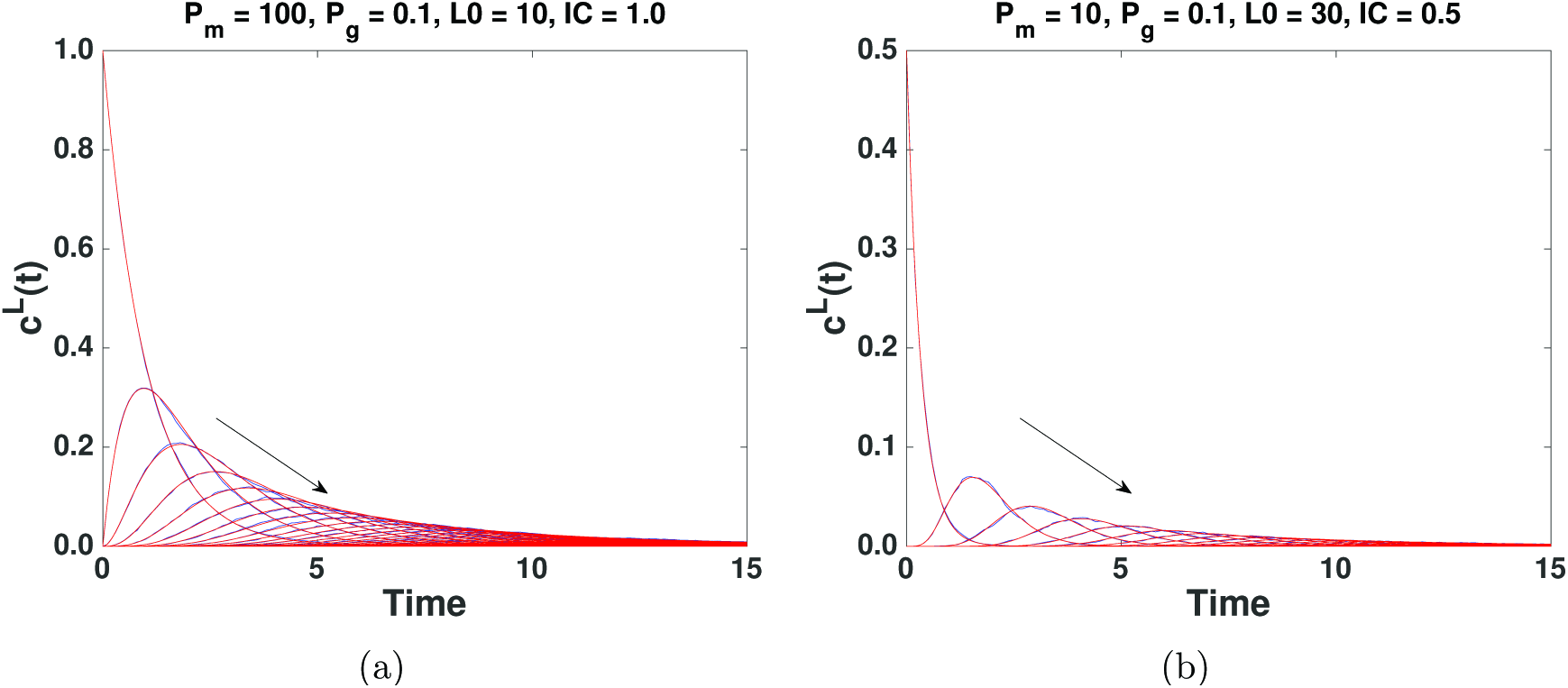
The evolution of the individual densities, *c*^*L*^*(t)*, for two different sets of parameters and initial conditions. *L*_0_ is the initial domain length, and IC is the initial density of agents. Numerical solution of Eq. (6) (red) and the corresponding ensemble averages from the IBM given by Eq. (13) (blue). In (a) domain lengths *L*_0_, *L*_0_ + 1,…, *L*_*trunc*_ are plotted, in (b) domain lengths *L*_0_, *L*_0_ + 5, …, *L*_*trunc*_ are plotted. Ensemble averages are taken from 10000 repeats. Increasing domain length is down and to the right in all panels, as indicated by the arrows.

In Fig. 3 the evolution of the pairwise densities, given by Eqs. (11) and (12), can be seen. In the top row of Fig. 3 the evolution for distances *r* = 1, 6 and 10 is displayed (panels (a)-(c) correspond to Fig. 2 (a)). We see an excellent agreement between the ensemble average of the IBM and the numerical solution of Eqs. (11) and (12) for all values of *r* as both time and domain length increases. In the bottom row of Fig. 3 the evolution for distances *r* = 1, 16 and 30 corresponding to Fig. 2 (b) is displayed. Again we see an excellent agreement for all values of r. It is important to note that the ensemble averages become increasingly noisy for larger values of r. This is because the number of samples in the ensemble average from the IBM decreases for each count, Eq. (17), as the variance in the domain length increases.

**Figure 3:**
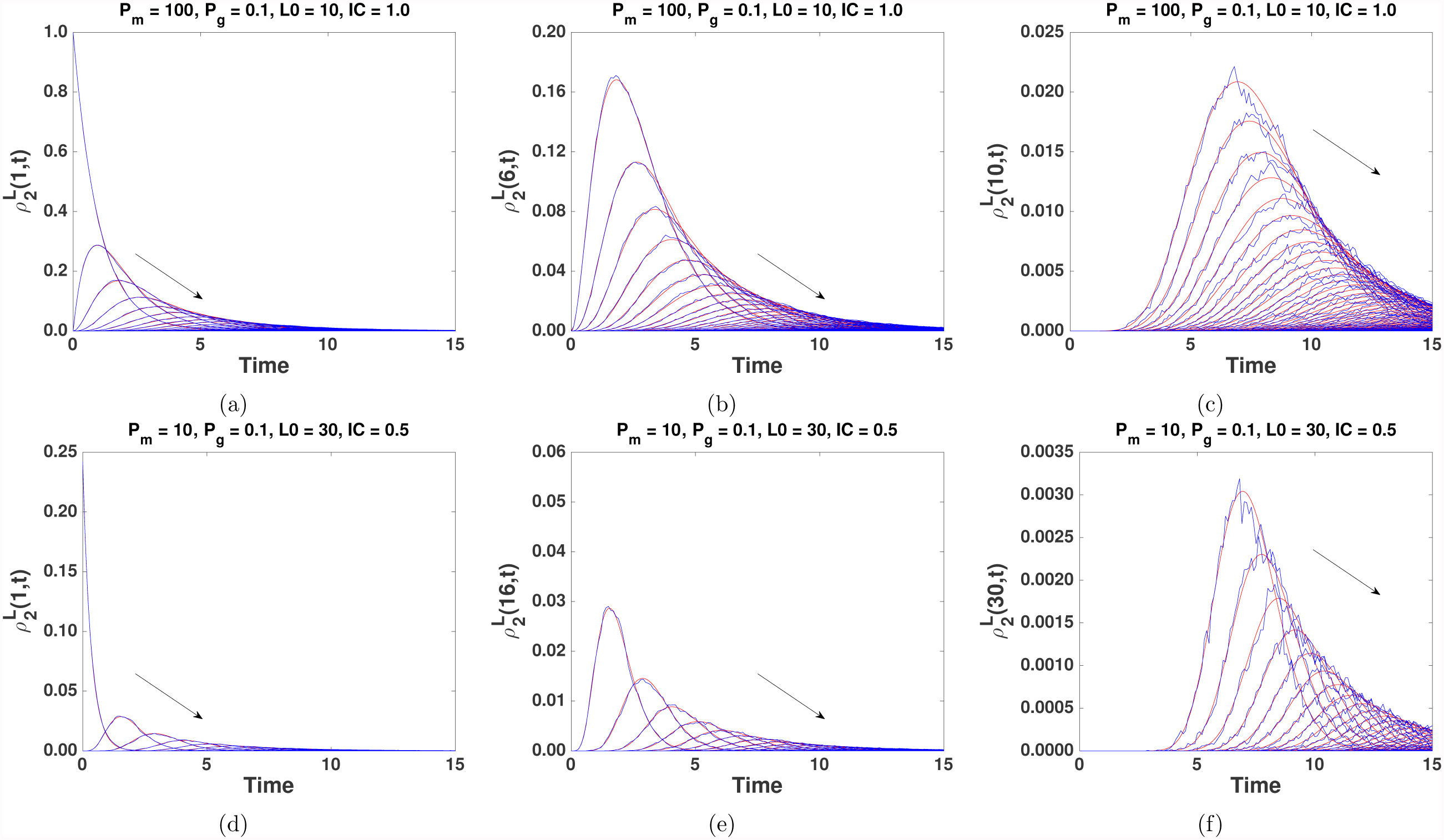
The evolution of the pairwise densities, 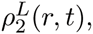, for two different sets of parameters and initial conditions (a)-(c) and (d)-(f) (panels (a)-(c) correspond to Fig. 2 (a) and (d)-(f) correspond to Fig. 2 (b)). Numerical solution of Eqs. (11) and (12) (red), and the corresponding ensemble averages from the IBM given by Eq. (17) (blue). Ensemble averages are taken from 10000 repeats. Top row: (a) The evolution of the pairwise density function at distance *r* = 1 for the domain length *L*_0_, *L*_0_ + 1,…, *L*_*trunc*_ Increasing domain length is down and to the right in all panels, as indicated by the arrows. (b) The evolution of the pairwise density function at distance *r* = 6. (c) The evolution of the pairwise density function at distance *r* = 10. Bottom row: (d) The evolution of the pairwise density function at distance *r* = 1 for the domain length *L*_0_, *L*_0_ + 5,…, *L*_trunc_. (e) The evolution of the pairwise density function at distance *r* = 16. (f) The evolution of the pairwise density function at distance *r* = 30.

### 3.2 Agent proliferation

We now add agent proliferation to Eqs. (11) and (12). The addition of agent proliferation will include three-point density terms in Eqs. (11) and (12), that is, the probability of three sites having a given occupancy. Similarly, an expression for three-point distribution terms would include four-point distribution terms and so on. Therefore, it is necessary to close the system of equations. To do so we employ the Kirkwood superposition approximation (KSA), which has been used successfully in scenarios where the domain does not grow [13, 16, 17]. The details of how to do this have been previously given and so we simply state the result, however, we provide them again in Appendix A. If we include agent proliferation in Eq. (6) we obtain

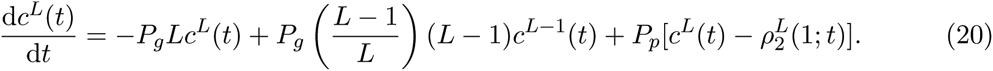

If we include agent proliferation in Eqs. (11) and (12), for the case where *r* = 1 we obtain

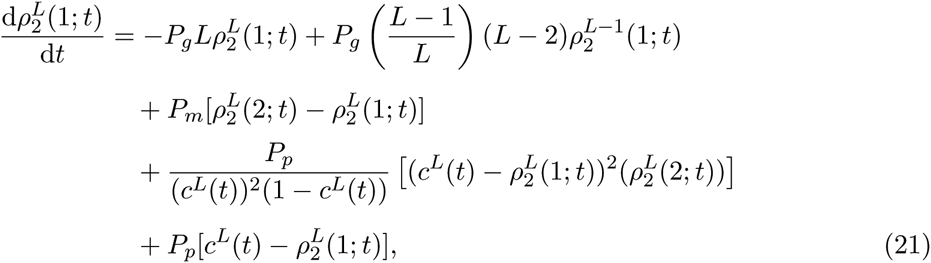

and for 1 < *r* < *L* we obtain

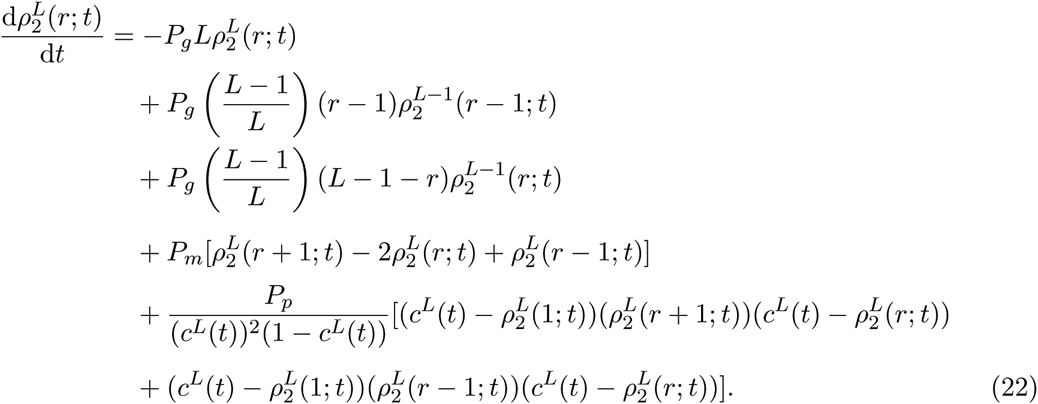

As can be seen from Eqs. (21) and (22) the inclusion of agent proliferation and associated implementation of the KSA means that individual density functions are now present in the equations for the evolution of the pairwise density functions, which is not the case without agent proliferation (Eqs. (11) and (12)). Similarly, the inclusion of agent proliferation means the evolution of the individual density functions now depends on the pairwise density functions, as can be seen in Eq. (20). It is important to note that Eqs. (21) and (22) are not exact due to the use of the KSA [13].

### 3.3 Motile, proliferative agents on a growing domain

We now present results for motile, proliferative agents on a growing domain. As before the initial condition in the IBM is achieved by populating a certain number of sites uniformly at random. We initialise the individual density functions corresponding to the initial domain length in the same manner as Eq. (15). However, to avoid a singularity in Eqs. (21) and (22) we follow Johnston et al. [18] and initialise all other individual densities with a small parameter ∊ = 10^−20^ [18]. That is,

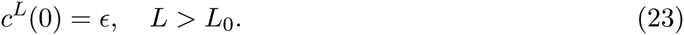

This small parameter was selected by iteratively reducing its size until the error tolerance threshold of the ODE numerical solver was compromised. This approach meant that the effect of introducing the small parameter ∊ into the equations was minimised.

In Fig. 4 the evolution of the individual density functions can be seen for two simulations with different parameters. In Fig. 4 (a) we see the evolution of the individual densities for a domain with an initial length of ten lattice sites and two agents. Unlike Eqs. (11) and (12), Eqs. (21) and (22) are not exact due to the use of the KSA. Despite this, we see that a good agreement is achieved between the numerical solution of Eq. (20) and the ensemble averages from the IBM for a low rate of agent proliferation. However, as we increase agent proliferation, relative to agent motility, the approximation by the numerical solution of Eq. (20) of the ensemble averages from the IBM becomes less accurate, as can be in Fig. 4 (b). This is because as agent proliferation is increased spatial correlations become more prevalent in the IBM [13, 16].

**Figure 4:**
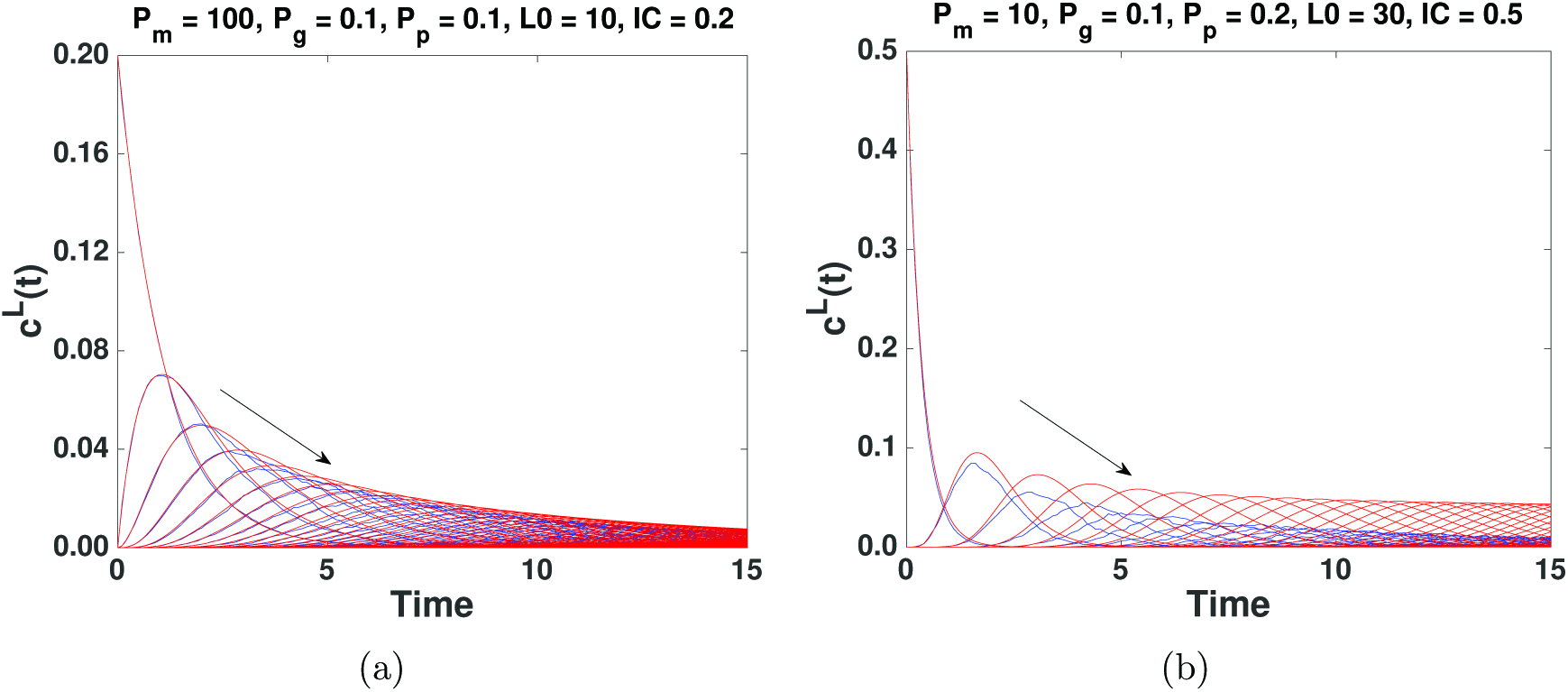
The evolution of the individual densities, *c*^*L*^*(t)*, with agent proliferation for two different sets of parameters and initial conditions. Numerical solution of Eq. (20) (red) and the corresponding ensemble averages from the IBM given by Eq. (13) (blue). In (a) domain lengths *L*_0_, *L*_0_+1, …, L_trunc_ are plotted, in (b) domain lengths *L*_0_, *L*_0_+5, …,L_trunc_ are plotted. Ensemble averages are taken from 10000 repeats. Increasing domain length is down and to the right in all panels as indicated by the arrows.

The error associated with the KSA also grows monotonically as time evolves, as can be seen in Fig. 4 (b) where the evolution of the individual density functions for a domain with an initial length of thirty lattice sites and fifteen agents is displayed. This error is for two reasons: i) the KSA introduces an error; and ii) the initialisation of the one-point distributions with a small parameter ∊ to avoid a singularity introduces an error.

In Fig. 5 the evolution of the pairwise densities can be seen. In the top row of Fig. 5, which corresponds to Fig. 4 (a), the evolution for distances *r* = 1, 6 and 10 is displayed. We see a good agreement for *r* = 1, 6 and 10. In the bottom row of Fig. 5, which corresponds to Fig. 4 (b), the evolution for distances *r* = 1, 16 and 30 is displayed. We initially see a good agreement for *r* = 1 and 16. This agreement begins to break down as time evolves for the two reasons discussed above.

**Figure 5:**
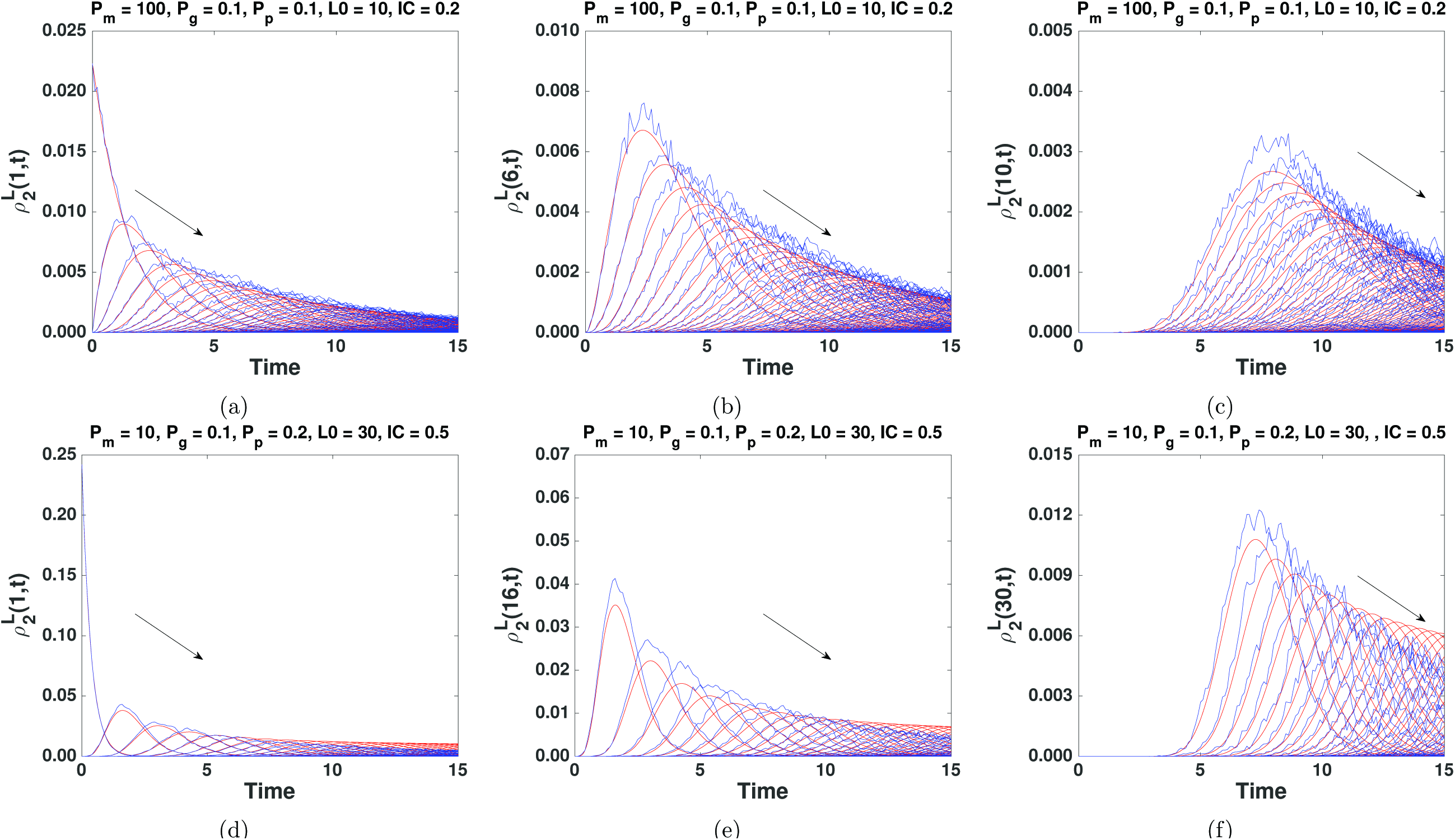
The evolution of the pairwise densities, 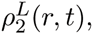 with agent proliferation for two different sets of parameters and initial conditions (a)-(c) and (d)-(f) (panels (a)-(c) correspond to Fig. 4 (a) and (d)-(f) correspond to Fig. 2 (b)). Comparison of the numerical solution of Eqs. (21) and (22) (red) and the corresponding ensemble averages from the IBM given by Eq. (17) (blue). Ensemble averages are taken from 10000 repeats. Top row: (a) The evolution of the pairwise density function at distance *r* = 1 for the domain length *L*_0_, *L*_0_ + 1; … ;*L*_trunc_, (b) at distance *r* = 6, and (c) at distance *r* = 10. Bottom row: (d) The evolution of the pairwise density function at distance *r* = 1 for the domain length *L*_0_, *L*_0_ + 5; …, *L*_*trunc*_, (e) at distance *r* = 16, and (f) at distance *r* = 30. Increasing domain length is down and to the right in all panels as indicated by the arrows.

## 4 Integrating domain growth into correlation functions

Our current framework, Eqs. (20)-(22), results in a system of individual density functions given by Eq. (20) which, while mathematically sound, is not easy to relate to biological systems and the experimental data associated with them, whereby the evolution of a macroscopic density on a growing domain is generally only measured with respect to time [3, 4, 8, 30]. Therefore, to mitigate these issues, we present a method to reduce the system of individual density functions given by Eq. (20) into a single equation describing the evolution of the macroscopic density on a growing domain. To do this we introduce ‘correlation functions’, and integrate the effect of domain growth into the pre-existing work on the calculation of spatial correlations between agents [13-18]. To derive this simplified model we will assume all domains are of length *L*, where *L* is the mean of the stochastic model.

To reduce the distribution of individual density functions given by Eq. (20) into a single equation describing the evolution of the macroscopic density on a growing domain we use the following heuristic approximations

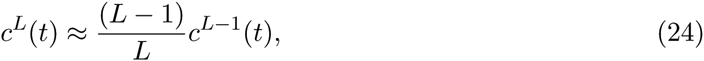

for the individual density functions, and

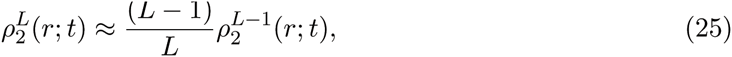

for the pairwise density functions. This is a previously published approximation [31] and implies that domain growth ‘dilutes’ both the agent density and the pairwise densities. To demonstrate the validity of Eqs. (24) and (25) we measure the relative error

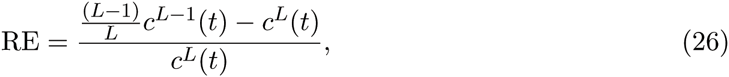

of approximations Eq. (24) and (25) and present the results in the Appendix B.

If we substitute approximation Eq. (25) into Eqs. (21) and (22) for *r* = 1 we obtain

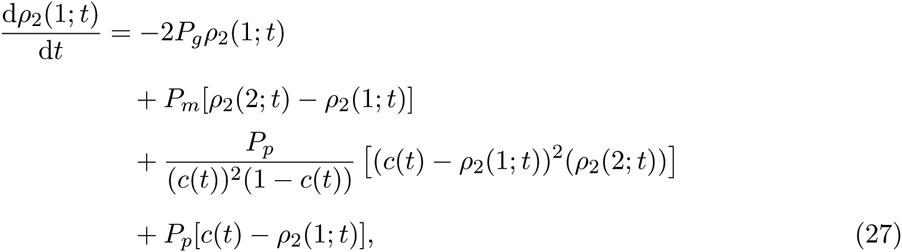

and for 1 < *r* < L

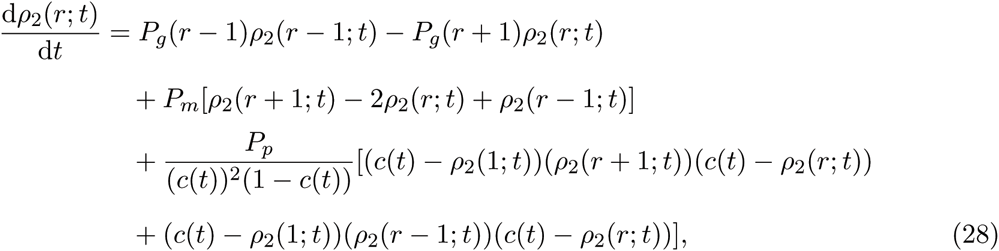

whereby all pairwise density functions are now for the same domain length, *L*, and so the superscript indicating domain length has been dropped. Similarly, when approximation Eq. (24) is applied to Eq. (20) it becomes

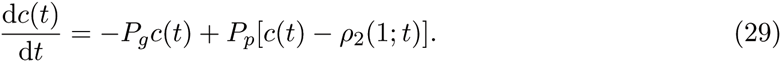

This is a single equation, albeit coupled, describing the evolution of the macroscopic density on a growing domain that includes the effect of spatial correlations between agents created by agent proliferation.

### 4.1 Correlation function

We finally introduce correlation functions. The correlation function [22, 23] is defined as

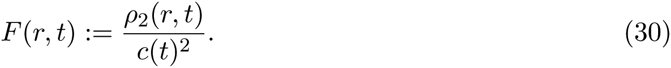

The correlation function is a useful change of variables to improve the visualisation of results, as *F (r, t)* ≡ 1 means that the occupancy of two lattice sites a given distance apart is independent. It has therefore been widely used in the study of spatial correlations in models of cell motility and proliferation [13-18].

Eq. (30) can now be substituted into Eqs. (27) and (28) to describe the evolution of spatial correlations between agents on a growing domain. This results in Eq. (27) becoming

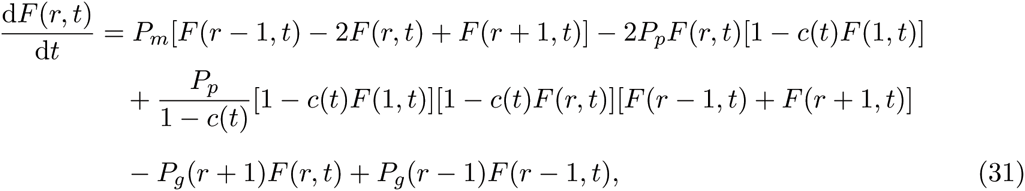

and Eq. (28) becoming

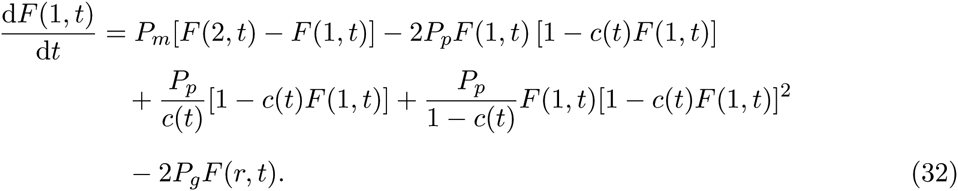

If we rewrite Eq. (29) in terms of Eq. (30) we obtain

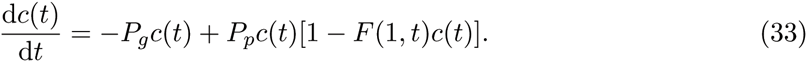

Eq. (33) is a hybrid (corrected) mean-field correlation ODE model (referred to from now on as the corrected ODE model), whereby the effect of agent proliferation on spatial correlations between agents is included in an equation describing the evolution of the agent density on a growing domain.

### 4.2 Results for correlation ODE model

We now compare simulations for motile, proliferative agents on a growing domain with the numerical solution of Eq. (33). As before the initial condition in the IBM is achieved by populating a certain number of sites uniformly at random. The initial length of the domain in all simulations is one hundred lattice sites. We also compare ensemble averages from the IBM with a *(uncorrected)* mean-field approximation (MFA)

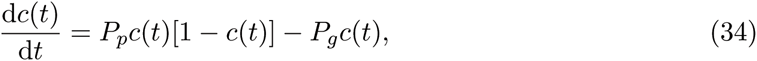

in which it has been assumed that *F*(1, *t*) = 1, i.e. the effect of spatial correlations is ignored. This means we compare ensemble averages from the IBM with both the MFA (Eq. (34)) and the corrected ODE model (Eq. (33)).

To solve Eqs. (31) and (32) numerically we use an implicit Euler scheme with the tridiagonal matrix algorithm and Picard linearisation. For all numerical solutions presented here *δt* = 0.1 and *Δx* = 0.1, and our correlation truncation value is *r* = 50. By this it is meant we set all correlation functions at and beyond this distance equal to one. That is,

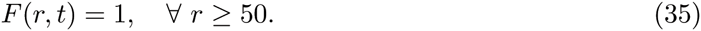

Similarly, our initial condition is that all distances are initially uncorrelated, that is

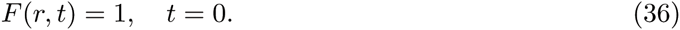

In Appendix C we provide results from simulations that involve only domain growth i.e. no agent proliferation or movement. However, we now turn our attention to simulations that include agent proliferation and motility. As we increase *P*_*p*_ relative to *P*_*g*_ the approximation of the spatial correlations in the IBM by the corrected ODE model becomes less accurate, as can be seen by comparing Fig. 6 (b) and (c), and Fig. 6 (e) and (f). This is due to both the KSA approximation and the use of Eqs. (24) and (25), and leads to the approximation of the evolution of the agent density in the IBM becoming less accurate as can be seen in Fig. 6 (a) and (d). However, in both scenarios the corrected ODE model more accurately approximates the evolution of the agent density in the IBM than the standard mean-field model Eq. (34).

**Figure 6:**
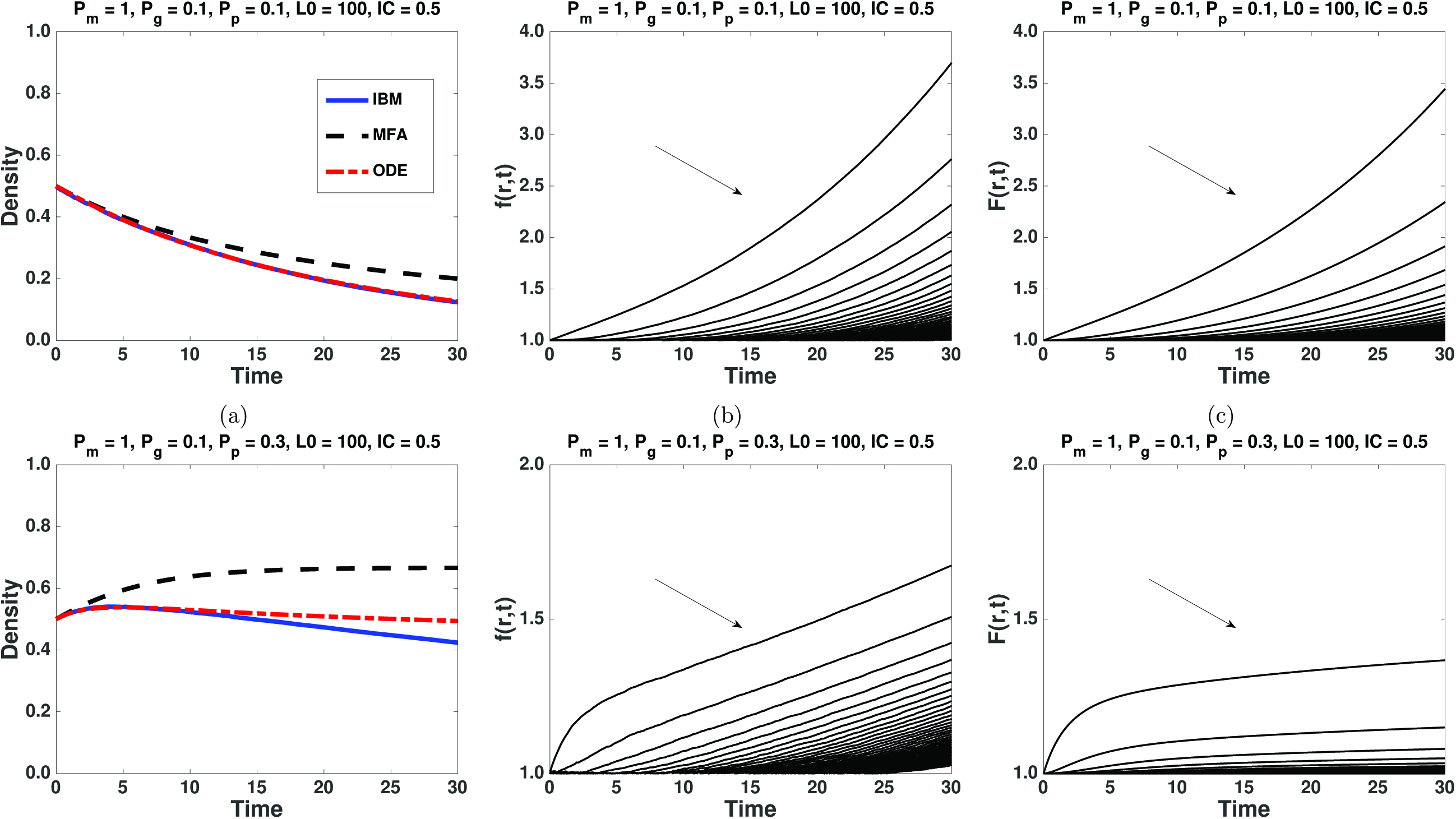
(Colour online): Panels (a) and (d) demonstrate the evolution of the macroscopic agent density with agent proliferation as given by Eqs. (13), (33) and (34). Panels (b) and (e) demonstrate the evolution of the pairwise correlations in the IBM. The correlation function is plotted for increasing distance from Δ to 50Δ in steps of Δ in the direction indicated by the arrow in panels (b), (c), (e) and (f). Panels (c) and (f) demonstrate the pairwise correlations at different distances from the numerical solution of Eqs. (31) and (32). The distance, *r*, increases from Δ to 50Δ in steps of Δ. The ensemble averages are taken from 100 repeats of the IBM.

## 5 Discussion

We have presented methods to include the effects of domain growth in the evolution of the individual and pairwise agent density functions in a one-dimensional IBM. Spatial correlations have been shown to play an important role in cell migration and tumour development, with both of these processes associated with domain growth [2-4, 8, 27]. For systems without agent proliferation comparisons between averaged discrete results and our corrected ODE model are excellent, as can be seen in Figs. 2 and 3. These results show that uniform domain growth, somewhat intuitively, reduces spatial correlations between agents. The introduction of agent proliferation into the correlation ODE model leads to some inaccuracy. However, there is still good agreement between the averaged discrete results and our corrected ODE model in parameter regimes with low proliferation as can be seen in Figs. 4 and 5. This inaccuracy is largely due to the use of the KSA approximation, and to a lesser extent the small parameter ∊ introduced into the model [13, 18].

Following this we integrated our framework into the existing literature on calculating the evolution of pairwise spatial correlations, namely, correcting mean-field approximations for the evolution of the agent density in the IBM [13-18]. This required a heuristic approximation [13], Eqs. (24) and (25), that allowed us to to derive a single equation describing the evolution of the macroscopic density in the IBM on a growing domain. This meant we could place the results naturally alongside the measurement of cell density on growing domains in experimental data [2-4, 24-26, 30]. Approximations Eqs. (24) and (25) were also necessary so that the effects of domain growth could be included in an ODE system describing the evolution of spatial correlations in a model of cell migration and proliferation. We demonstrated the accuracy of this ODE formulation in Figs. 6 and 8 (Appendix C).

The results presented here are extendable to non-periodic boundary conditions and systems where we do not assume translational invariance. In addition, the framework presented could also be extended to higher-dimensional models of cell motility and proliferation.

## Appendix A A.1 Individual density function

On a growing domain the individual density functions for motile and proliferative agents evolve according to

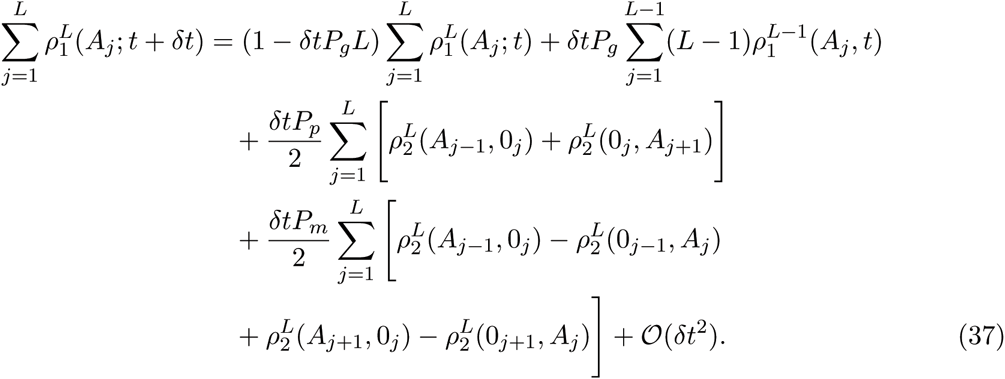

If we assume translational invariance we have 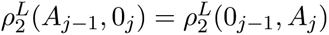 and 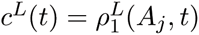 = 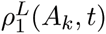 and, recognising that 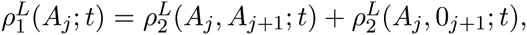 we can simplify Eq. (37) to obtain

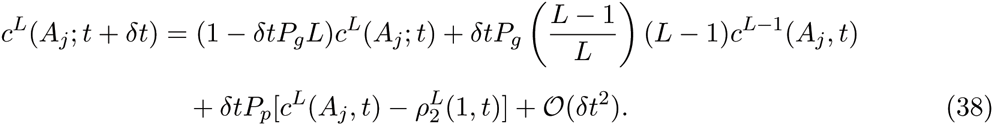

If we rearrange Eq. (38) and take the limit as *δt* −> 0 we arrive at

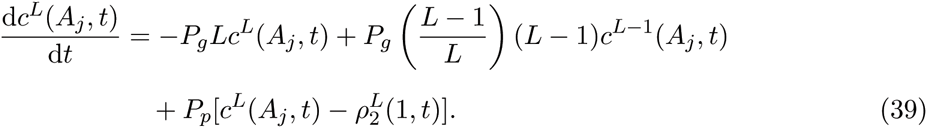

**A.2 Pairwise density functions**

On a growing domain the pairwise density functions for motile and proliferative agents for the distance *r* = 1 evolve according to

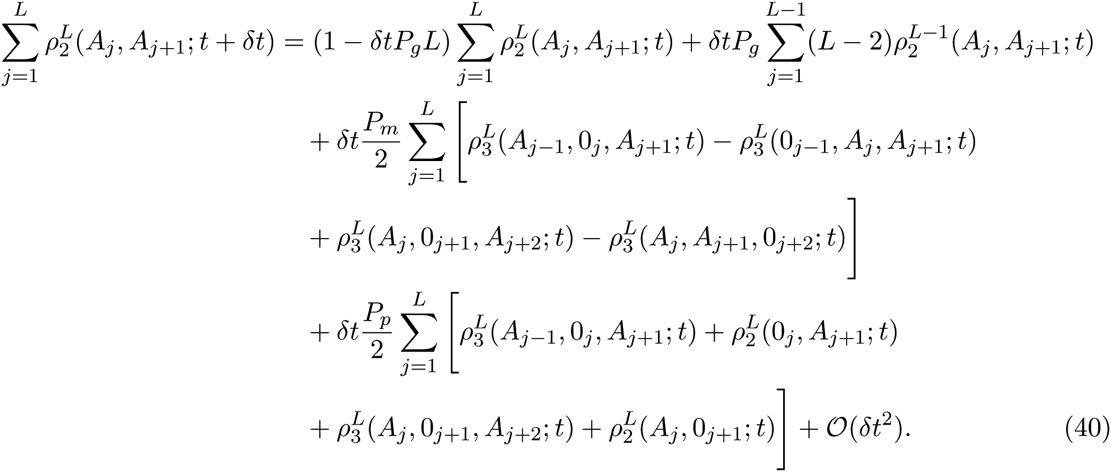

If we assume translational invariance and recognise that

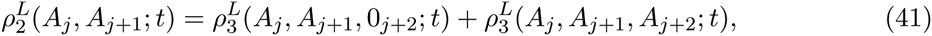

and

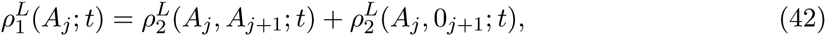

we obtain

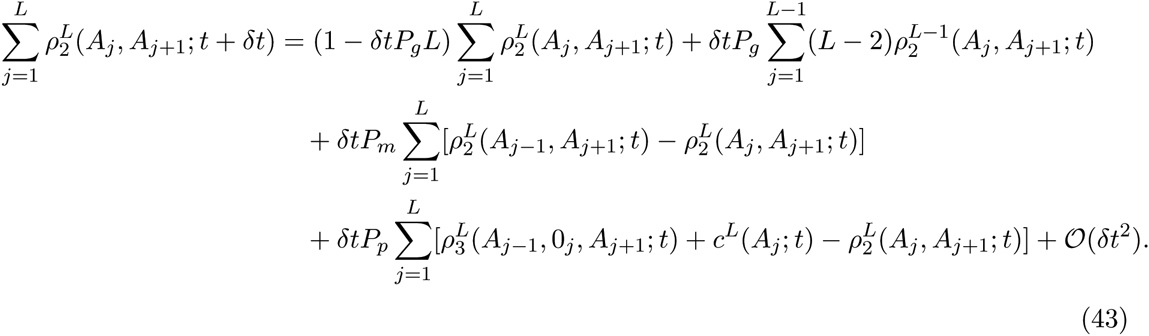

If we now implement the Kirkwood superposition approximation [13] (this allows us to write three-point density functions in terms of pairwise density functions, and so close our system of equations Eqs. (39) and (43)), that is

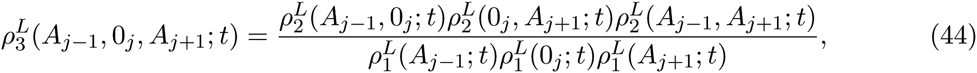

we arrive at

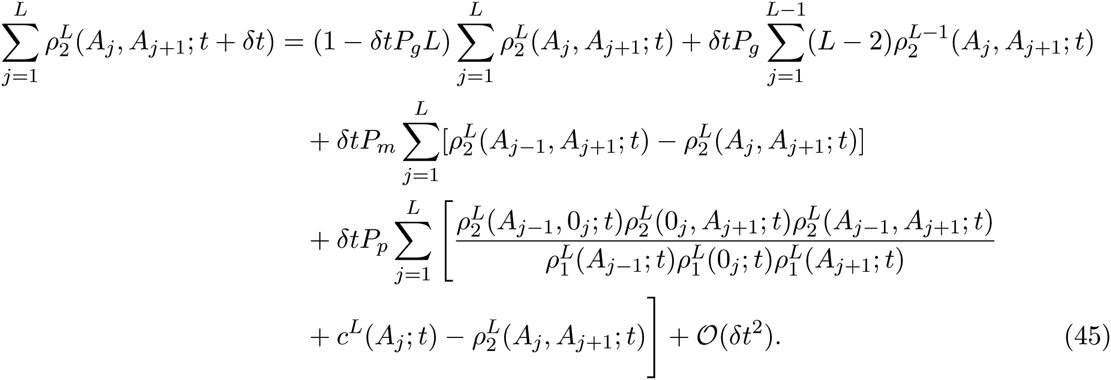

Eq. (45) can be simplified to obtain

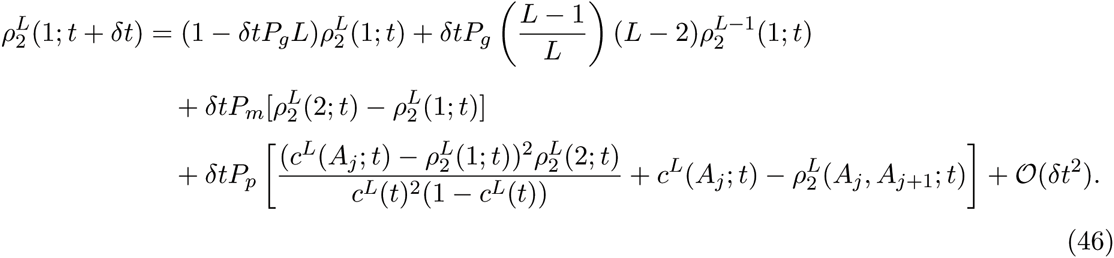

If we take the limit as *δt* → 0 Eq. (46) becomes

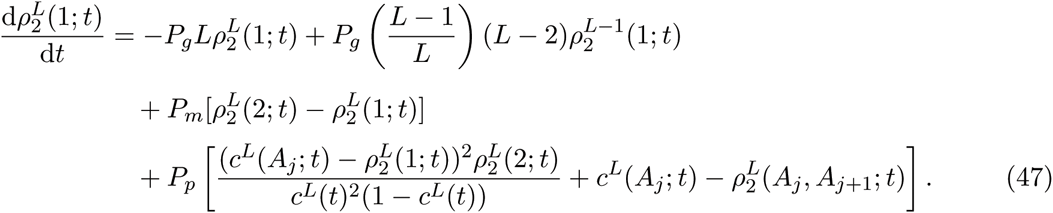

Eq. (47) is just Eq. (21) in the main text.

On a growing domain the pairwise density functions for motile and proliferative agents for distances 1 < *r* < *L* evolve according to

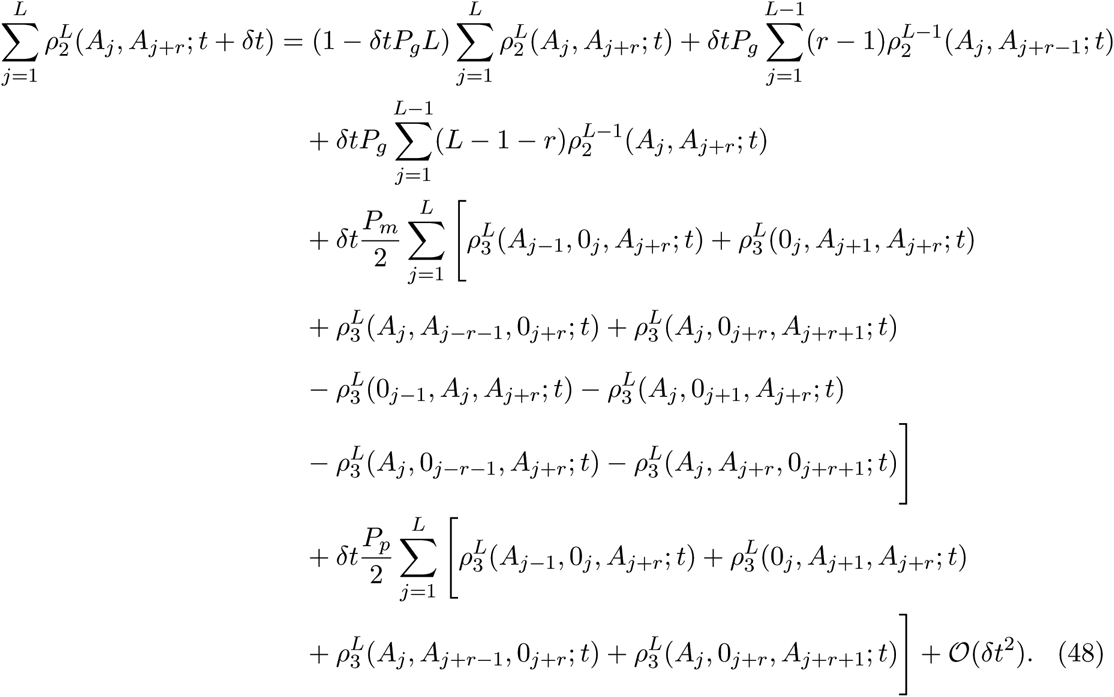

From Eq. (48) we can obtain Eq. (22).

## Appendix B

In Fig. 7 the error caused by approximation Eq. (24) for the individual density functions for two different simulations can be seen. The relative error is initially high, due to the initial conditions (Eqs. (15) and (23)), but decreases towards zero as the expected domain length increases. The nature of the error caused by the approximation Eq. (25) for the pairwise density functions is the similar to the individual density functions and so is not shown.

**Figure 7:**
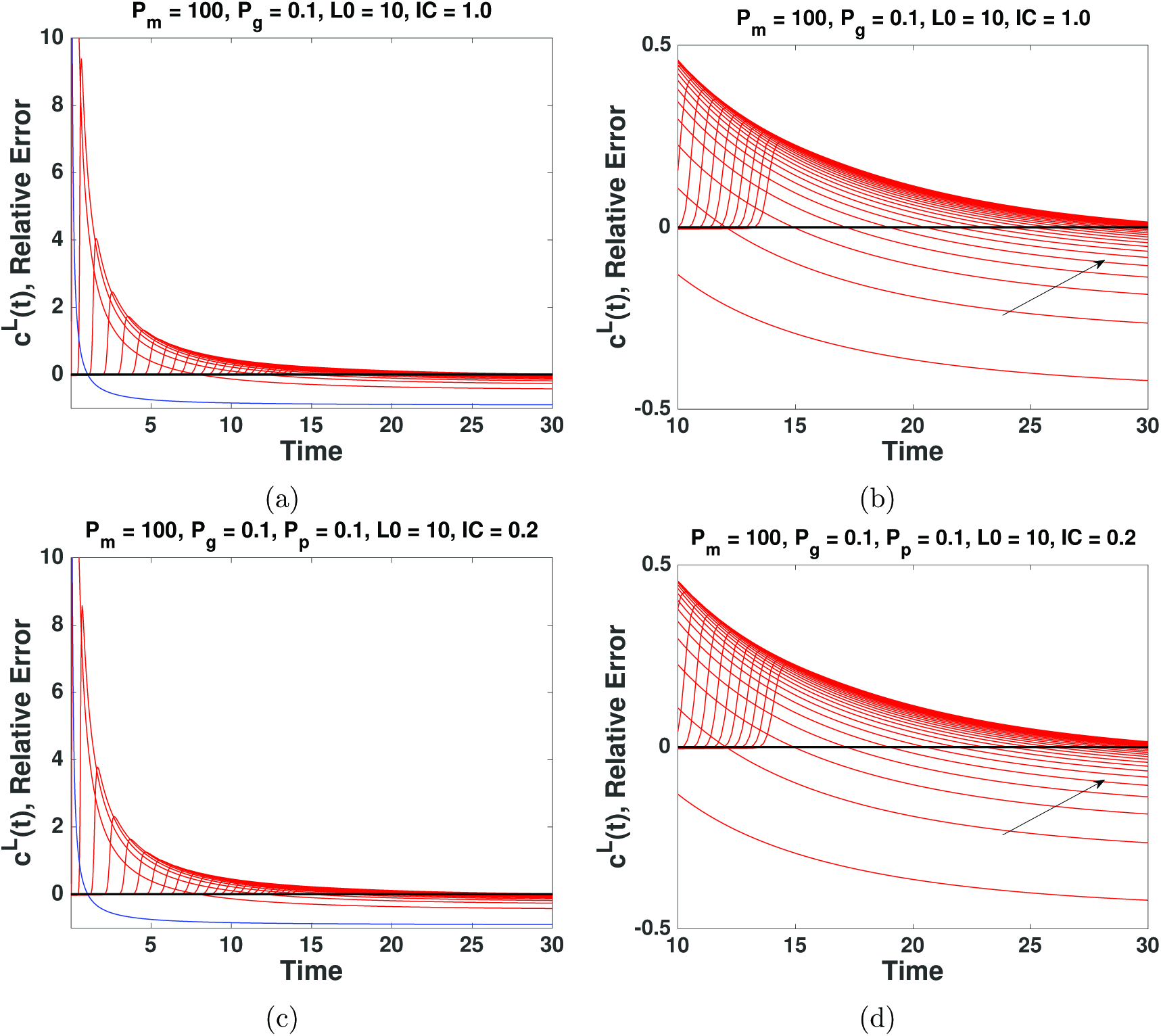
The relative error, calculated with Eq. (26), in the individual densities due to approximations Eq. (24) and (25). The domain length increases in distances of 10Δstarting from the initial domain length *L*_0_ (indicated by the blue line). In panels (a) and (c) increasing domain length is up and to the right of the blue line. Panels (b) and (d) are magnified aspects of panels (a) and (c), and increasing domain length is indicated by the black arrows. We can see that the relative error approaches zero as the domain length increases. The black line indicates a relative error of zero.

## Appendix C

In Fig. 8 (b) we see the effects of exponential domain growth on spatial correlations between agents in the IBM. Domain growth does not create spatial correlations between agents that are initially distributed uniformly at random (when averaged over many repeats). This is indicated by the correlation function remaining at a value of one, and is also captured by the numerical solutions of Eqs. (31) and (32), as can be seen in Fig. 8 (c) and (f). As the position of the agents remains uncorrelated throughout the course of the simulation, the evolution of the macroscopic density in the IBM, mean-field and corrected ODE models all match exactly as can be seen in Fig. 8 (a) and (d).

**Figure 8:**
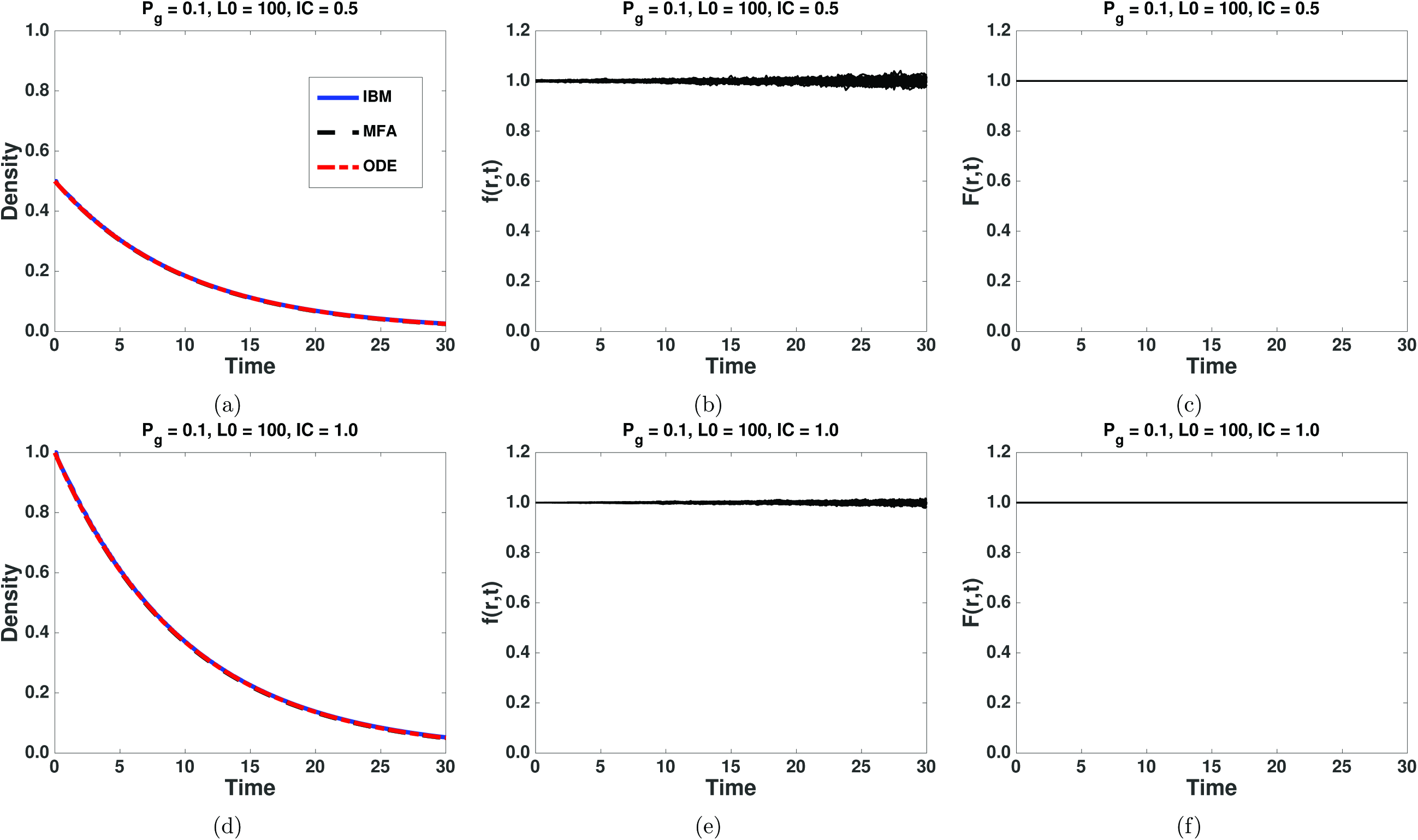
(Colour online): Panels (a) and (d) demonstrate the evolution of the macroscopic agent density as given by Eqs. (13), (33) and (34). Panels (b) and (e) demonstrate the evolution of the pairwise correlations in the IBM. The correlation function is plotted for increasing distance from Δto 50Δin steps of Δin panels (b) and (e). Panel (c) and (f) demonstrate the pairwise correlations from the numerical solution of Eqs. (31) and (32). The distance, *r*, increases from Δto 50Δin steps of Δ. The ensemble averages are taken from 100 repeats of the IBM.

